# Discovery of a novel monoclonal PD-L1 antibody H1A that promotes T-cell mediated tumor killing activity in renal cell carcinoma

**DOI:** 10.1101/2022.09.10.507426

**Authors:** Zesheng An, Michelle A. Hsu, Joanina K. Gicobi, Tianxiao Xu, Susan M. Harrington, Henan Zhang, Kevin D. Pavelko, Jacob B. Hirdler, Christine M. Lohse, Reza Nabavizadeh, Rodriguo R. Pessoa, Vidit Sharma, R. Houston Thompson, Bradley C. Leibovich, Haidong Dong, Fabrice Lucien

**Affiliations:** Department of Immunology, Mayo Clinic, Rochester, MN, USA; Department of Urology, Mayo Clinic, Rochester, MN, USA; Department of Laboratory Medicine and Pathology, Mayo Clinic, Rochester, MN, USA; Immune Monitoring Core, Mayo Clinic, Rochester, Minnesota; Division of Clinical Trials and Biostatistics, Department of Quantitative Health Sciences, Mayo Clinic, Rochester, Minnesota

**Keywords:** immune checkpoint blockade, PD-L1, renal cell carcinoma, adjuvant therapy, individualized medicine

## Abstract

In the last decade, the therapeutic landscape of renal cell carcinoma has rapidly evolved with the addition of PD-1/PD-L1 immune checkpoint inhibitors in the armamentarium of oncologists. Despite clinical evidence of improved oncological outcomes, only a minority of patients experience long-lasting antitumor immune response and complete response. The intrinsic and acquired resistance to PD-1/PD-L1 immune checkpoint blockade is an important challenge for patients and clinicians as no reliable tool has been developed to predict individualized response to immunotherapy. In this study, we demonstrate the translational relevance of an ex-vivo functional assay that measure the tumor cell killing ability of patient-derived CD8 T cells isolated from peripheral blood. Cytotoxic activity of CD8 T cells was improved at 3-month post-radical nephrectomy compared to baseline and it was associated with higher circulating levels of tumor-reactive effector CD8 T cells (CD11a^high^CX3CR1^+^GZMB^+^). Pretreatment of peripheral immune cells with FDA-approved PD-1/PD-L1 inhibitors enhanced tumor cell killing activity of CD8 T cells but differential response was observed at the individual patient level. Finally, we found a newly developed monoclonal antibody (H1A), which induces PD-L1 degradation, demonstrated superior efficacy in promoting T-cell mediated tumor killing activity compared to FDA-approved PD-1/PD-L1 inhibitors. PBMC immunophenotyping by mass cytometry revealed enrichment of effector CD8 T cells in H1A-treated PBMC. To conclude, our study lays the ground for future investigation of the therapeutic value of H1A as a next-generation immune checkpoint inhibitor. Furthermore, further work is needed to evaluate the potential of measuring T-cell cytotoxicity activity as a tool to predict individual response to immune checkpoint inhibitors in patients with advanced renal cell carcinoma.

## Introduction

Renal cell carcinoma (RCC) is amongst the top 10 most common cancers in both men and women with an estimated 75,000 cases each year in the US. The prognosis is poor with 30% of patients presenting with metastatic disease at diagnosis and approximately 30% of those who initially present with localized disease develop metastases post-surgery. Immune checkpoint inhibitors (ICIs) have revolutionized the therapeutic landscape of patients with advanced RCC. PD-1/PD-L1 checkpoint blockade has become front-line therapy with four different treatments approved by the FDA since 2018[1]. The high rates of disease recurrence post-surgery in patients with RCC has led to the testing of immune checkpoint inhibitors in the adjuvant setting. In a phase III study (KEYNOTE-564 trial), the efficacy of pembrolizumab (Keytruda, Merck), a humanized anti-PD-1 antibody, was investigated as adjuvant treatment for RCC patients with high-risk of disease recurrence post-nephrectomy[2]. Following clinical observation of improved disease-free survival compared to placebo, pembrolizumab has been approved by the FDA in 2021, becoming the first immunotherapy for adjuvant treatment of RCC. It remains that only a minority of patients demonstrate complete or durable response to immune checkpoint inhibitors. At 30-month follow-up, 25% of patients treated with pembrolizumab developed local or distant recurrence compared to 35% for the placebo group at 2-year which indicates the need for patient selection. Two other phase III trials (Checkmate-914, Immotion010) evaluating adjuvant nivolumab (anti-PD-1) plus ipilimumab (anti-CTLA-4) (Opdivo plus Yervoy, BMS) and atezolizumab (anti-PD-L1) (Tecentriq, Roche) did not meet their primary endpoint of disease-free survival but a cohort of carefully selected patients may benefit from these agents.

Expression of PD-L1 in primary RCC tumors has been extensively investigated as potential predictive biomarker of response to PD-1/PD-L1 inhibitors[3]. In past clinical trials, metastatic RCC patients with PD-L1-positive tumors showed better objective response rates to PD-1/PD-L1 blockade compared to patients with PD-L1 negative tumors. However, several limitations hinder the use of PD-L1 status to guide treatment decision such as PD-L1 expression discordance between primary tumors and metastases[4], variability in PD-L1 assays and scoring algorithms[5]. Furthermore, in the recent KEYNOTE-564 trial, no significant difference was seen in response to adjuvant pembrolizumab in patients with PD-L1-positive and - negative tumors[2]. With the advent of immuno-oncology drugs in the adjuvant setting of RCC, there is an urgent need to develop tools to select the patients who will benefit the most from a particular immunotherapeutic regimen.

The paradoxical response to PD-1/PD-L1 inhibitors in patients with PD-L1-negative tumors and inversely suggest that PD-L1 expression in non-tumor cells may also contribute to tumor immune escape, hence, be better predictor of response to immunotherapy. PD-L1 expression has been shown in myeloid cells[6] and antibody blockade of PD-L1 in antigen presenting cells is essential for tumor control in preclinical models[7, 8]. Based on this evidence, we sought to assess the impact of blocking immune cell-derived PD-L1 signaling on T-cell mediated tumor cell killing. Herein, we isolated peripheral blood mononuclear cells from patients surgically treated for RCC and quantify T-cell mediated tumor cell killing in response to FDA-approved PD-1/PD-L1 inhibitors. Similarly, we assessed for the first time the therapeutic potential of a newly developed monoclonal anti-PD-L1 antibody (H1A clone) that destabilizes PD-L1 on cell surface and induces its degradation[9].

## Materials and Methods

### Blood collection and patient characteristics

Blood was collected from 14 patients undergoing radical nephrectomy for clear-cell renal cell carcinoma (ccRCC). Demographics and clinicopathological characteristics can be found in **Table 1**. All samples were collected prior to surgery (baseline) and at 3-month post-surgery with approved Mayo Clinic Institutional Review Board (IRB #16-006956). Whole patient blood was drawn in EDTA-coated vacutainers (BD Biosciences, San Jose, CA). Peripheral blood mononuclear cells (PBMC) were isolated from patient blood by centrifugation with Lymphoprep (StemCell Technologies, Vancouver, Canada) and SepMate devices (StemCell Technologies, Vancouver, Canada). PBMC were resuspended in Cryostor freeze medium and stored in liquid nitrogen until use. Platelet-poor plasma was obtained from centrifugation of whole blood at 2,500 x g for 15 minutes at room temperature. Plasma was aliquoted in cryovials and stored at -80°C until use.

**Table 1.** Demographics and Patient Characteristics.

|  | Number of Patients | % |
| --- | --- | --- |
|  | 14 | 100 |
| **Age (years)** |  |  |
| Mean (range) | 62 (39-74) |  |
| **Gender** |  |  |
| Male | 8 | 57 |
| Female | 7 | 43 |
| **T Stage** |  |  |
| T1a | 3 | 21.4 |
| T1b | 2 | 14.2 |
| T2b | 1 | 7.1 |
| T3a | 4 | 28.5 |
| T3b | 3 | 21.4 |
| T4 | 1 | 7.1 |
| **N Stage** |  |  |
| N0 | 7 | 50.0 |
| N1 | 1 | 7.1 |
| Nx | 6 | 42.9 |
| **M Stage** |  |  |
| M0 | 8 | 57.1 |
| M1 | 6 | 42.9 |
| **Tumor Grade** |  |  |
| 2 | 5 | 35.7 |
| 3 | 4 | 28.6 |
| 4 | 5 | 35.7 |
| **Tumor Necrosis** |  |  |
| No | 4 | 28.6 |
| Yes | 10 | 71.4 |
| **Tumor Size (cm)** |  |  |
| Mean (range) | 8.1 (1.7-13.4) |  |

### PBMC activation and treatment

Human PBMC were thawed, washed with Hank’s Balanced Salt Solution (HBSS) twice and resuspended in RPMI 1640 (Corning, New York, NY, USA) supplemented with 10% heat inactivated FBS (Thermo Fisher Scientific, Waltham, MA), rhIL2 (10 U/ml), rhIL15 (10 ng/ml) and rhIL7 (10 ng/ml) (Peprotech, Cranbury, NJ). PBMC were incubated overnight followed by addition of soluble anti-CD3 (5 μg/mL; catalog no. 555336, BD Pharmingen) to allow PBMC activation prior to T-cell mediated tumor cell killing assay. For treatment with anti–PD-1/PD-L1 immune checkpoint inhibitors, antibodies were added at the time of activation and at 20 μg/ml.

### Cell lines

Human 786-O ccRCC cell line (ATCC, Manassas, VA, USA) was maintained in a humidified incubator at 5% CO_2_. 786-O cells were cultured in RPMI 1640 (Corning, New York, NY, USA) with 10% fetal bovine serum (Thermo Fisher Scientific, Waltham, MA) and 5% penicillin and streptomycin (Thermo Fisher Scientific, Waltham, MA). PD-L1 was knocked-out in 786-O cells using CRISPR/Cas9 technology to exclude any direct effect of anti-PD-L1 antibody on tumor cells in the tumor killing assay.

### T-cell mediated tumor cell killing

Target 786-O cells were trypsinized and labeled with Calcein-AM (5 μmol/L, catalog no. C1430, Invitrogen) for 30 minutes at 37°C in dark, washed with Hank’s Balanced Salt Solution (HBSS) twice and resuspended at 1 × 10^5^/mL in RPMI 1640 medium. Human PBMC and target cells were mixed at target to effector ratio 1:20 in FBS-free RPMI 1640 medium and seeded into wells of a 96-well U-bottom plate. Plate was briefly centrifuged at 300 x g for 1 minute, then incubated at 37°C for 4 hours. Saponin (0.1%) was added to wells that contained only tumor cells to provide a value for “maximum calcein release.”. In addition, target cells without effector cells were used to measure spontaneous release of calcein and medium only was used to measure fluorescence background. All experimental and control conditions were performed in four replicates. After 4-hour incubation, the plate was centrifuged at 500 x g for 5 minutes. Hundred microliters of the 200-μL supernatant was transferred to a new, opaque (black) 96-well flat-bottom plate (catalog no. 237105, Thermo Fisher Scientific). Calcein fluorescence was read using an automated fluorescence measurement system (BioTeK Synergy HTX multi-mode reader) with an excitation of 485 × 20 and an emission filter of 528 × 25, scanning for 1 second per well. Read-outs were exported to Microsoft Excel. Values for fluorescence background (medium only) were subtracted to all experimental and control values. To calculate the percentage of cytotoxicity, we used the following formula: % cytotoxicity = 100 x [(B-A)/(C-A)] with A=values for spontaneous release, B=values for experimental conditions and C=values for maximum release).

### Soluble PD-L1 ELISA

For detection of soluble PD-L1 in blood of RCC patients, ELISA plates (Immulon 2HB 96-well plates, Biolegend) were coated with 100 ng per well (100 μl) of monoclonal antibody against PD-L1 (H1A clone) overnight at 4 °C. Free binding sites were blocked with 200 μl of blocking buffer (Pierce) for 1 hr at room temperature. Then 50 μl of plasma samples were mixed with BSA 1% and added to each well in duplicates. After overnight incubation at 4°C, biotinylated monoclonal PD-L1 antibody (B11 clone) was added to each well and incubated for 1 hr at room temperature. A total of 100 μl per well of horseradish peroxidase-conjugated streptavidin (BD Biosciences) diluted in PBS containing 0.1% BSA was then added and incubated for 1 hr at room temperature. Plates were developed with tetramethylbenzidine (Pierce) and stopped with 0.5N H_2_SO_4_. The plates were read at 450 nm with a BioTek plate reader. Recombinant human PD-L1 protein (R&D Systems, Cat# 156-B7) was used to make the standard curve ranging from 0.3125 to 10 ng/ml.

### Mass Cytometry by time-of-flight (CyTOF)

Human PBMC from five RCC patients were treated with PBS, atezolizumab and H1A for 72 hours as described above. Cells were counted on a Countess II automated cell counter and approximately 1×10^6^ cells were prepared for antibody labeling. Cells were blocked with 1 μg anti-mouse Fc block (BD Biosciences) for 10 minutes at room temperature followed by staining with a cocktail of 36 metal-conjugated antibodies for the human lymphocyte panel (**Supplementary Figure 1B**). The antibodies were obtained from Fluidigm or generated in-house by the Mayo Clinic Hybridoma Core. Surface marker staining was performed in cell staining buffers (Fluidigm) at room temperature for 30 minutes. Intracellular staining was performed using a Cytofix/Cytoperm Kit (BD Biosciences) per the manufacturer’s protocol. DNA intercalation was performed by adding 1:10000 diluted 125 μM of Cell-ID™ Intercalator-Ir (Fluidigm) with gentle agitation in 4°C for 30 minutes. Upon completion of staining, cells were stored in fresh 1% methanol-free formaldehyde in PBS (Thermo Fisher Scientific) until the day of data collection. All events were acquired on a Helios mass cytometer (Fluidigm). Randomization, bead normalization, and bead removal of the data collected were performed using CyTOF software (Fluidigm).R-based tool Cytofkit was used for initial gating of ^191^Ir and ^193^Ir DNA intercalators as well as the event length parameter to discern intact singlets from debris and cell aggregates. Live/dead staining was then used to identify live intact singlets. All CD45^+^ cells were gated and used for subsequent t-SNE and CITRUS analysis. Clustering and dimensionality reduction to 20,000 events per file was performed using the Rphenograph algorithm (t-distributed stochastic neighbor embedding) that included all 36 markers in the panel. For t-SNE analysis, files were categorized into either PBS-treated, atezolizumab-treated or H1A-treated (n=5 per group). Individual and cleaned FCS files were exported for analysis and uploaded to the Cytobank software platform (Beckman Coulter) for subsequent analysis with CITRUS (cluster identification, characterization, and regression) algorithm. CITRUS identified clusters of cell subpopulations that are statistically associated or correlated with an experimental phenotype of interest (responder/non-responder). CITRUS was performed using the Significance Analysis of Microarrays (SAM) correlative association model (Benjamini-Hochberg-corrected P value at a False Discovery Rate of 1%) with the following parameters: Clustering channels: “select all 36 antibody-metal channels”; Compensation: “File-Internal Compensation”; Association Models: “Significance Analysis of Microarrays (SAM)—Correlative”; Cluster Characterization: “Abundance”; Event sampling: “Equal”; Events sampled per file: “3,701”; Minimum cluster size (%): “2”; Cross Validation Folds: “5”; False Discovery Rate (%): “1”. Once maps were generated, the relative expression level of each marker of interest could be visualized.

### Statistical Analysis

Student *t*-test (parametric), Mann-Whitney test (non-parametric) was employed to compare two groups. One-way ANOVA (parametric) and Kruskal-Wallis (non-parametric) tests were used to compare three or more groups. The results were considered significant for p values <0.05. p values were either specified in the figure or denoted as asterisk: * p<0.05, ** p<0.01, *** p<0.001, **** p<0.0001. All data were analyzed and plotted in GraphPad Prism 9.3.0.

## Results

### Improved tumor cell killing by peripheral T cells following radical nephrectomy

While regarded as an immunogenic tumor, RCC elicits immune dysfunction through infiltration of immunosuppressive cells and expression of immuno-inhibitory molecules including PD-L1[10, 11]. We sought to determine the impact of radical nephrectomy on peripheral CD8 T cell function using our *ex-vivo* platform to quantify the cytotoxic activity of CD8 T cells[12]. Five out of 11 samples (45.4%) showed detectable killing at baseline (pre-nephrectomy) (**Figure 1A**). The median killing rate of patient-derived CD8 T cells was 11.4% with a range from 4.0% to 32.5%. At 3-month post-nephrectomy, 72.7% (7/11 samples) showed killing activity with 6 patients showing an elevation in killing efficiency and 1 patient showing a decrease from baseline to 3-month post nephrectomy (11.4% vs 0%). Four patients showed no T-cell killing efficiency at both baseline and 3-month post-surgery. Median killing rate at 3-month post-nephrectomy was 21.4% (range: 0%-52.5%), 1.9 times higher than at baseline (p=0.0245). In our cohort, 4 out of 11 patients (36.3%) had metastatic disease at the time of radical nephrectomy. By subgrouping patients based on the metastasis status, no significant difference was observed in the killing rates of CD8 T cells at baseline and post-nephrectomy (**Figure 1B**). Median tumor cell killing rates were also not significantly different between non metastatic and metastatic RCC at both baseline and post-RN. Our ex-vivo data demonstrate that radical nephrectomy is associated with enhanced tumor cell killing ability of peripheral CD8 T cells even in patients who present with metastatic disease at diagnosis.

**Figure 1.**
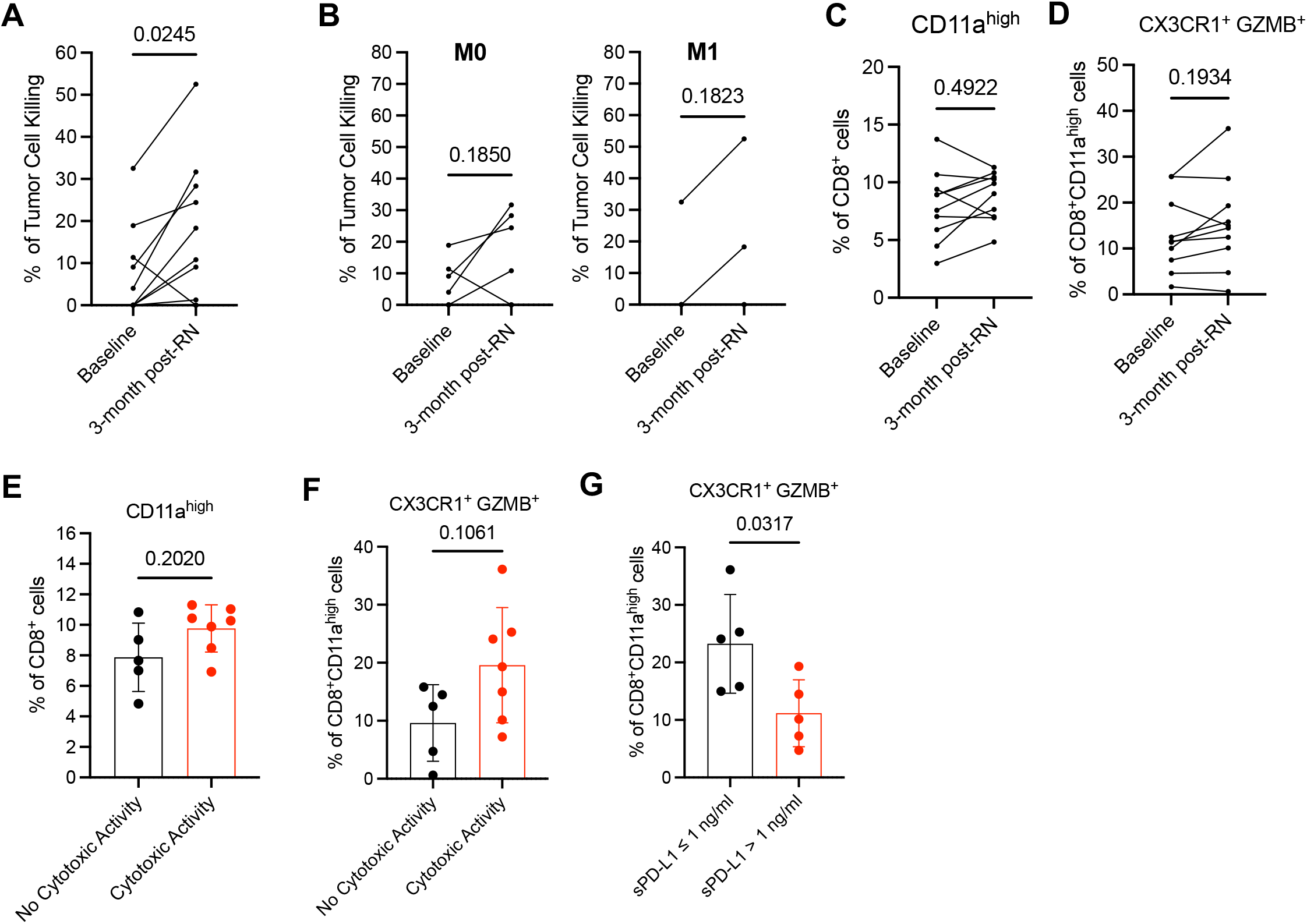
Improved tumor cell killing activity by peripheral T cells following radical nephrectomy. A) Percentage of tumor cell killing by peripheral CD8 T cells isolated from RCC patients prior to (baseline) and 3-month post radical nephrectomy (RN). N= 11 patients, Wilcoxon paired t test. B) Percentage of tumor cell killing by peripheral CD8 T cells isolated from RCC patients stratified by metastasis status at diagnosis, M0= no evidence of metastatic disease (N=7 patients), M1= evidence of metastatic disease (N= 4 patients), Wilcoxon paired t test. C) Percentage of peripheral tumor-reactive CD8^+^ cells defined as positive for CD11a^high^ in patients at baseline and 3-month post-RN. Wilcoxon paired t test. D) Percentage of peripheral tumor-reactive CD11a^high^CD8^+^ cells positive for effector markers CX3CR1 and GZMB in patients at baseline and 3-month post-RN. Wilcoxon paired t test. E) Percentage of peripheral tumor-reactive CD8^+^ cells defined as positive for CD11a^high^ in patients subgrouped by detectable ex vivo cytotoxic activity. Mann Whitney test. Data are presented in mean +/-sd. F) Percentage of peripheral tumor-reactive CD11a^high^CD8^+^ cells positive for effector markers CX3CR1 and GZMB in patients subgrouped by detectable ex vivo cytotoxic activity. Mann Whitney test. Data are presented in mean +/-sd. G) Percentage of peripheral tumor-reactive CD11a^high^CD8^+^ cells positive for effector markers CX3CR1 and GZMB in patients subgrouped by blood levels of soluble PD-L1. Mann Whitney test. Data are presented in mean +/-sd.

To explain the enhanced killing activity of peripheral CD8 T cells following radical nephrectomy, we measured blood levels of tumor-reactive CD11a^high^CD8 T cells expressing CX3C chemokine receptor 1 (CX3CR1) and Granzyme B (GZMB). We and others have shown that CD11a is highly expressed in peripheral CD8 T cells in response to tumor antigen presentation[13, 14]. A subset of recently activated tumor-reactive CD11a^high^CD8 T cells expresses CX3CR1 and GZMB and show improved cytotoxicity activity leading to better patient outcome [14-18]. Levels of tumor-reactive CD11a^high^CD8 T cells post-RN were increased in 7 out of 11 patients (% increase = 44%) compared to baseline (**Figure 1C**). Only 2 patients showed decreased levels of CD11a^high^CD8 T cells and 2 patients had unchanged levels. For CD11a^high^CD8 T cells positive for CX3CR1 and GZMB, 6 out of 11 patients showed increased levels post-RN, 2 patients decreased levels and 3 patients unchanged levels (**Figure 1D**).

Both ex vivo cytotoxicity assay and CD8 T cell immunophenotyping was performed in PBMC isolated from 12 patients at 3-month post-nephrectomy (**Figure 1E-1F**). Five samples showed no evidence of cytotoxic activity while 7 samples showed detectable activity with a median tumor cell killing of 26.4% (range: 10.8%-52.5%). The percentage of CD8 T cells expressing high levels of CD11a was slightly higher in patients showing cytotoxic activity (mean 10.27% vs 7.7%, p=0.20) (**Figure 1E)**. Cytotoxic activity of CD8 T cells was associated with higher levels in tumor-reactive CD8 T cells positive for the effector markers CX3CR1 and GZMB (median 9.62% vs 19.6%, p=0.11) (**Figure 1F)**. Differential levels of circulating tumor-reactive effector T cells in patients may result from the presence of immunosuppressive signals preventing expansion of peripheral tumor-reactive CD8 T-cells. In the form of soluble protein or carried by extracellular vesicles, PD-L1 has been recently showed to suppress T-cell activity locally and systemically[19-21]. Furthermore, high concentrations of circulating PD-L1 have been associated to worse prognosis and resistance to immune checkpoint blockade [21-23]. In our study, we observed lower levels of tumor-reactive effector CD8 T cells in patients with blood concentrations of PD-L1 superior to 1 ng/ml (median=6.146 ng/ml) compared to patients with less than 1 ng/ml of circulating PD-L1 (median=0.121 ng/ml) (p=0.032) (**Figure 1G**).

These findings demonstrate that peripheral expansion of tumor-reactive CD8 T cells with an effector phenotype is a prerequisite for effective cytotoxic killing of tumor cells. Furthermore, our data supports circulating PD-L1 as a negative regulator of systemic antitumor immune response in RCC patients.

### Treatment of patient-derived PBMCs with anti-PD-1/PD-L1 immune checkpoint inhibitors enhanced CD8 T tumor cell killing activity

Targeting PD-L1/PD-1 interaction between tumor and immune cells has been the primary focus for the development of immune checkpoint inhibitors. Recent studies highlighted the important role of PD-L1 expressed by myeloid cells in modulating tumor-specific CD8 T cell activation and function[6-8]. Here, we tested the efficacy of FDA-approved immune checkpoint inhibitors to stimulate the cytotoxic function of patient-derived CD8 T cells collected at 3-month post nephrectomy. To avoid interference with PD-L1 expressed by tumor cells, we used 786-0 tumor cells knocked out for PD-L1 as target cells. In PBMC treated with pembrolizumab (anti-PD-1), 10 out of 11 samples showed increased in T-cell killing rates with a median of 37.8% (range: 17.4%-57.7%) in pembrolizumab group compared to 14.6% (range: 0%-52.5%) in the control group (**Figure 2A**). No detectable killing activity was found in 6 samples of the control group but upon pembrolizumab treatment, tumor cell killing was observed in all samples. At the individual level, median increase of killing rates between control and pembrolizumab-treated cells was 18.3% (range: 0%-44.4%). In the nivolumab-treated group (anti-PD-1), all samples responded to treatment and similar increase to pembrolizumab group was observed with a median killing rate of 36.6% (range: 22.7%-65.4%) (**Figure 2B**). In the atezolizumab-treated group (anti-PD-L1), one patient showed no difference in T-cell cytotoxic activity compared to control (**Figure 2C**). Median killing rate in the atezolizumab group was 29.9% (range: 10.0%-52.5%) with a median increase of 13.8% compared to control group. Individual response to immune checkpoint inhibitors is heterogeneous with some patients showed stronger response with nivolumab while others had better killing activity with pembrolizumab (**Figure 2D**). Overall, no best response was observed with atezolizumab across all patients. Higher response with nivolumab was observed in 9 patients but it was not statistically significant. Three patients showed higher response to pembrolizumab and 1 patient had no difference in killing rates between both treatments. Altogether, our data demonstrate that treatment of PBMC with immune checkpoint inhibitors enhance T-cell tumor cell killing function but individual response varies with drug treatment.

**Figure 2.**
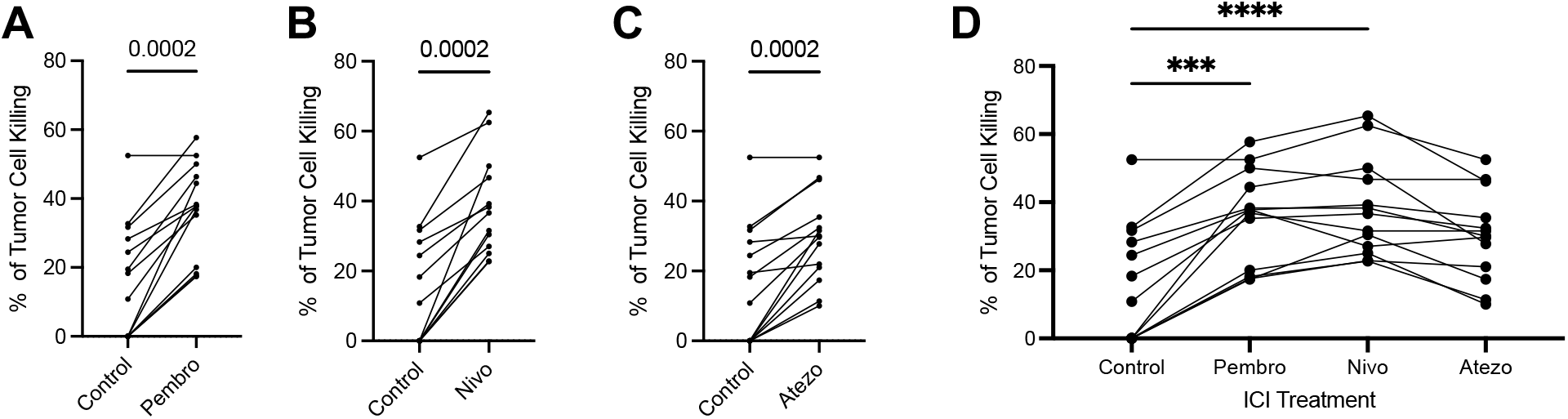
Treatment of patient-derived PBMCs with anti-PD-1/PD-L1 immune checkpoint inhibitors enhances CD8 T tumor cell killing activity. A) B) C) Percentage of tumor cell killing by peripheral CD8 T cells following patient-derived PBMC treatment with pembrolizumab (A), nivolumab (B), atezolizumab (C). N= 14 patients, Wilcoxon paired t test. D) Comparison of tumor cell killing rates between treatment groups and control group. Dunn’s multiple comparison test.

### Discovery of H1A, a novel monoclonal PD-L1 antibody that outcompetes FDA-approved immune checkpoint inhibitors

Our group has recently produced a novel monoclonal antibody that induces PD-L1 destabilization at the cell surface resulting in its degradation[9]. In tumor cells, we demonstrated that H1A-mediated PD-L1 degradation increases sensitivity to radiotherapy and DNA-damaging agents. Here, we sought to evaluate the efficacy of H1A in promoting a T-cell mediated tumor cell killing from RCC patient-derived PBMC. All patients responded to H1A treatment compared to control group (**Figure 3A**). Median killing rate in H1A-treated group was 41.7% (range: 25.0%-71.2%) which corresponds to an increase of 28.3% compared to control group. Overall, median killing rate was significantly higher in the H1A group than in the pembrolizumab group (p=0.0003) and the nivolumab group (p=0.003) (**Figure 3B**). In terms of individual response, all H1A-treated patients showed improved tumor cell killing efficiency compared to other treatment groups.

**Figure 3.**
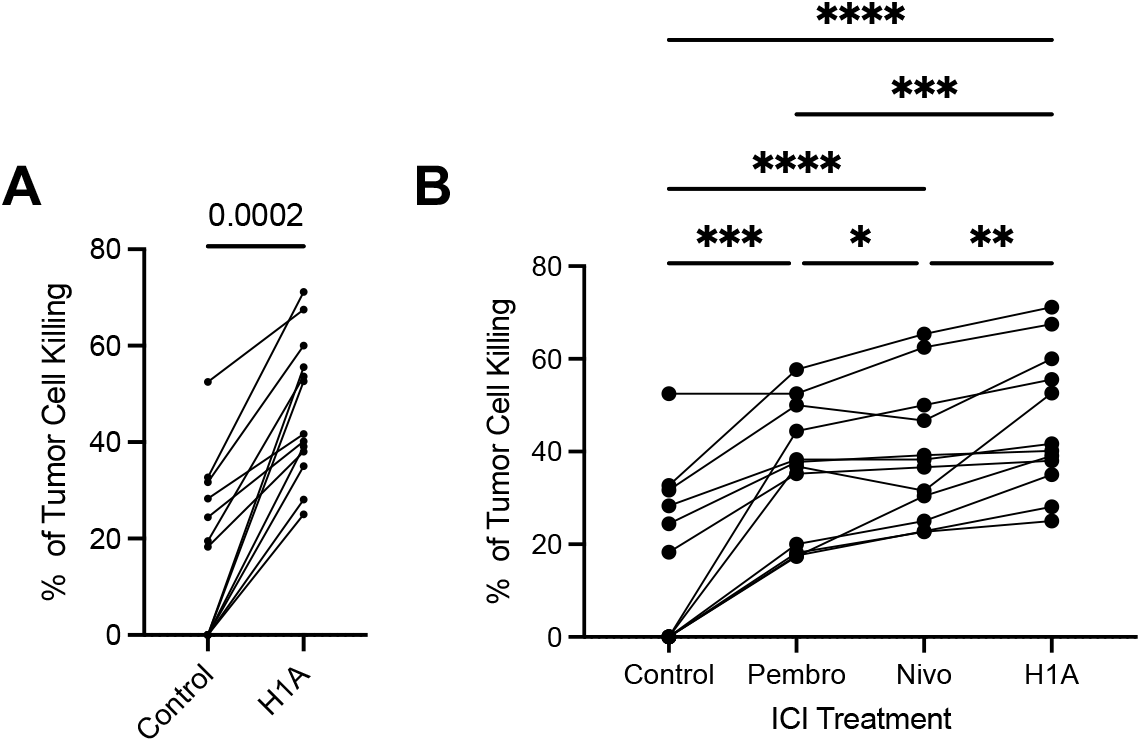
Discovery of H1A, a novel monoclonal PD-L1 antibody that outcompetes FDA-approved immune checkpoint inhibitors. A) Percentage of tumor cell killing by peripheral CD8 T cells following patient-derived PBMC treatment with anti-PD-L1 antibody clone H1A. N= 14 patients, Wilcoxon paired t test. B) Comparison of tumor cell killing rates between treatment groups and control group. Dunn’s multiple comparison test.

### Single-cell mass cytometry reveals enrichment in effector CD8 T cells in patient-derived PBMC treated with H1A

Our data demonstrate that, despite they both target PD-L1, atezolizumab and H1A have distinct functional impact on the tumor cell killing activity of CD8 T cells. Therefore, we investigated the differences in the immune cell composition and phenotype of PBMC treated with PBS (negative control), atezolizumab and H1A using single-cell mass cytometry. We selected PBMC from 5 patients who showed improved tumor cell killing in response to H1A (+ 40.5%, range: +34.1%-+55.6%, compared with PBS) while no significant difference was observed with atezolizumab (+14.2%, range: +2.4%-+27.8% compared with PBS). Immune cell abundance profiles of PBMC activated with soluble CD3 and treated with PBS and PD-L1 antibodies were analyzed using dimensionality reduction and t-SNE maps (**Figure 4A**). CD4 T cells, CD8 T cells were identified as the most abundant immune cell lineages and NK cells (CD4^-^CD8^-^CD16^+^) and B-cells (CD19^+^) as minor populations. Using RphenoGraph clustering algorithm, 26 clusters were identified which represent immune cell subpopulations such as regulatory T cells (CD25^+^,Foxp3^+^), effector T and NK cells (Granzyme B^+^, NKG7^+^) (**Figure S1A**). Differential expression of inhibitory immune checkpoint molecules such as PD-1, LAG-3 and TIGIT was found in both CD4 and CD8 T cells (**Figure S1A-S1B)**. T-SNE maps of PBS-, atezolizumab-(ATZ), and H1A-treated PBMC showed several clusters with differential abundance between groups (**Figure 4B**). Hierarchical clustering of Phenograph clusters showed that ATZ-treated PBMC are closer to PBS-treated samples than H1A-treated samples (**Figure 4C**). Three out of five H1A-treated samples showed high association in cluster enrichment which suggests similar PBMC phenotypes following H1A treatment. Several PBMC clusters were differentially and significantly enriched in response to ATZ and H1A treatment (**Figure 4D**). Five clusters (4, 5, 8, 18, 19) were more abundant in H1A-treated samples while five other clusters (1, 14, 16, 19, 21) were enriched in ATZ-treated samples. We employed CITRUS algorithm to identify the signature of immune cell subsets statistically associated with response to ATZ and H1A treatment. CITRUS performs an unsupervised hierarchical clustering to identify clusters of cellular populations within the overall dataset. This produced a map of nodes (a “tree”) with parent nodes upstream of their relative progeny (**Figure 4E**). Then, the correlative method SAM is used to discover stratifying signatures from clustered data features with statistical significance. Among CD4 T cells, the node #36965 and its progeny #36942 were significantly more abundant in ATZ-treated PBMC compared to H1A-treated samples (FDR 1%) (**Figure 4F**). Overlay of the relative expression of various markers onto the CITRUS map revealed high expression of the marker of regulatory T cells Foxp3 and the marker of proliferation Ki67 in these two clusters (**Figure 4G**). In line with this, ATZ-treated PBMC were enriched in proliferative regulatory T cells compared to H1A-treated PBMC (**Figure 4H**). Clusters #36965 and #36942 were also enriched in regulatory T cells positive for the inhibitory immune checkpoint molecules PD-1 and LAG-3 following ATZ treatment (**Figure 4I-4J**). In the CD8 T cell population, the cell cluster #36994 was enriched in H1A-treated samples and it was defined by high expression of Granzyme B (GZMB) and T-bet, two drivers of effector CD8 T cells (**Figure 4K**). H1A-treated PBMC were significantly enriched in GZMB^+^ T-bet^+^ CD8 T cells compared to ATZ-treated samples (**Figure 4L**). Similarly, a significant increase in NK cells positive for GZMB and Ki67 was observed in the H1A group compared to ATZ (**Figure 4M)**. Altogether, our mass cytometry-based immunophenotyping demonstrate that ATZ treatment leads to enrichment in immunosuppressive regulatory T cells while H1A treatment promotes expansion of effector CD8 T and NK cells.

**Figure 4.**
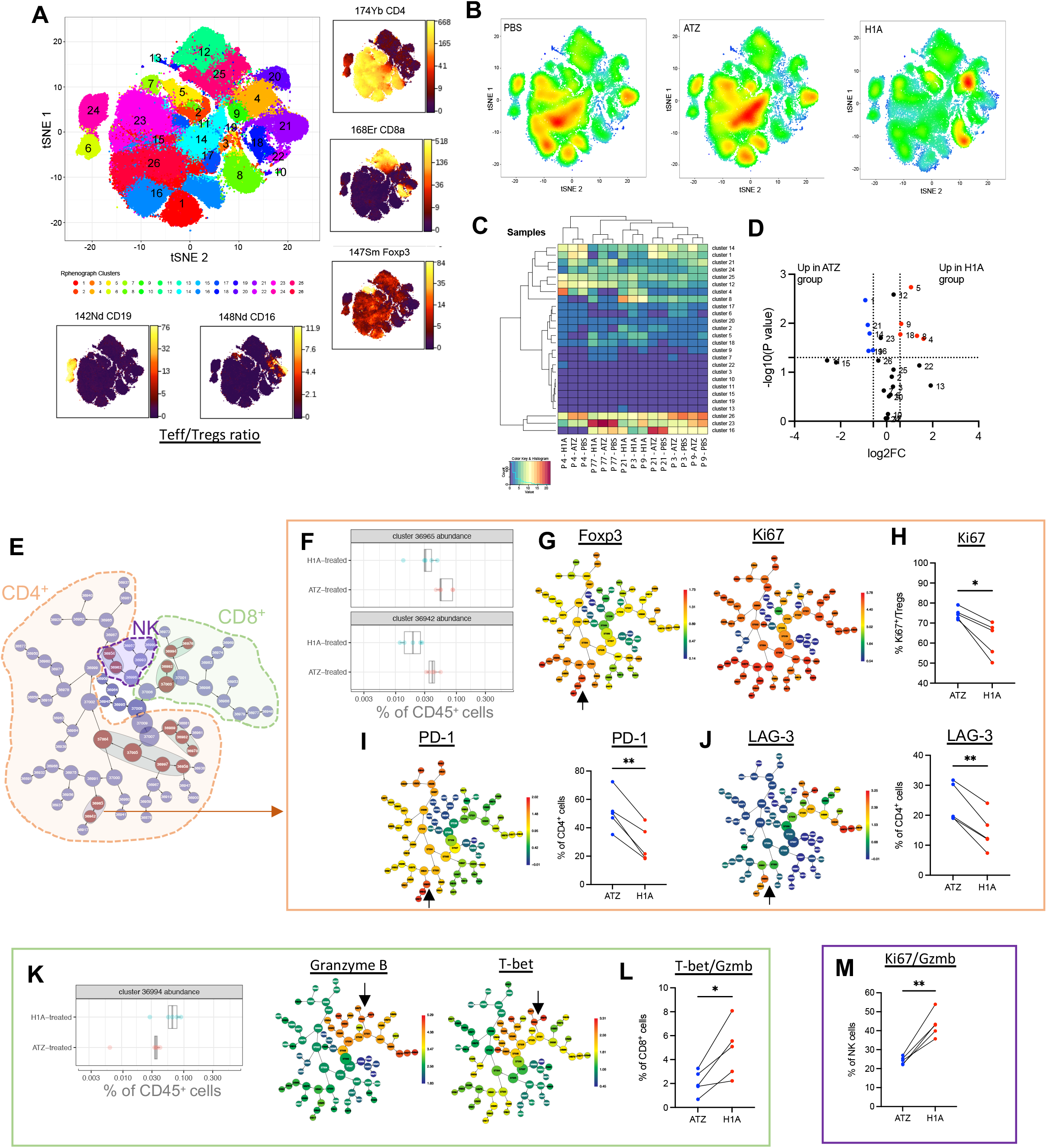
Single-cell mass cytometry reveals enrichment in effector CD8 T cells in patient-derived PBMC treated with H1A. A) T-SNE plot showing 26 cell clusters defined by the expression of 36 immune cell markers in RCC patient-derived PBMC (N=15 samples). Density of major immune cell markers (CD4, CD8, Foxp3, CD19 and CD16) are presented. B) Representative tSNE plots of each experimental group (PBS, atezolizumab (ATZ), H1A). Red indicates high-frequency categorization of cells to a cluster; blue indicates low frequency. C) Hierarchical clustering showing abundance of cell clusters in each patient treated with PBS, ATZ and H1A. D) Volcano plot showing cell clusters significantly upregulated in ATZ-treated samples (blue) and H1A-treated samples (red) (FC>1.5, adj p value<0.05). E) CITRUS-based clustering showing significant difference in abundance of cell populations of CD45^+^ cells treated with ATZ and H1A (N=5 patients, SAM correlative method with FDR 1%). F) Cluster abundance for #36942 and #36965 (CD4^+^ subset) shown as boxplot (n=5 patients, CITRUS with SAM and FDR 1%). G) Density of Foxp3 and Ki67 in the CITRUS clusters. Arrow indicate clusters #36942 and #36965. H) Percentage of Ki67+ regulatory cells (CD4^+^CD25^+^Foxp3^+^) in H1A- and ATZ-treated PBMC (N=5 patients). Paired t-test, *p<0.05. I) J) Density of PD-1 and LAG-3 in the CITRUS clusters and percentage of CD4^+^ T cells positive for PD-1 and LAG-3 in H1A- and ATZ-treated PBMC (N=5 patients). Paired t-test, **p<0.01. K) Cluster abundance for #36994 (CD8^+^ subset) shown as boxplot (n=5 patients, CITRUS with SAM and FDR 1%). Density of Granzyme B and T-bet in the CITRUS clusters. L) Percentage of GZMB+ T-bet^+^ CD8^+^ T cells in H1A- and ATZ-treated PBMC (N=5 patients). Paired t-test, *p<0.05. M) Percentage of GZMB+ Ki67^+^ NK cells in H1A- and ATZ-treated PBMC (N=5 patients). Paired t-test, **p<0.01.

## Discussion

In this study, we investigated for the first time the efficacy of PD-1/PD-L1 immunotherapies in eliciting patient-derived T-cell cytotoxic activity. Our study stands out by being the first to determine the functional consequences of targeting PD-L1/PD-1 signaling in the peripheral immune compartment. We employed our *ex vivo* T-cell tumor cell killing assay to measure the ability of patient-derived CD8 T cells to kill tumor cells in response immune checkpoint blockade. We also tested a newly generated monoclonal antibody (H1A) inducing cell-surface PD-L1 degradation[9]. Our study demonstrates the utility of an ex-vivo T-cell cytotoxicity assay to quickly assess response to immune checkpoint blockade in an individualized manner. Furthermore, it provides a scientific justification to dissect the mechanism(s) of action of immune checkpoint blockade in peripheral immune cells and its impact on antitumor immunity. Finally, our work lays the ground for future studies evaluating the therapeutic potential of H1A as next-generation adjuvant therapy for RCC.

The clinical benefit of radical (cytoreductive) nephrectomy in patients with metastatic RCC treated with immune checkpoint inhibitors is object of debate. The use of upfront cytoreductive nephrectomy is justified by the biological rationale that debulking of the primary tumor mass may eliminate immuno-inhibitory signals released by tumor cells and suppressor immune cells[24, 25]. Prior to the immune checkpoint inhibitor era, CARMENA and SURTIME trials showed that immediate cytoreductive nephrectomy does not improve clinical outcome in metastatic RCC patients treated with tyrosine kinase inhibitors[26, 27]. Survival benefit of upfront cytoreductive nephrectomy prior to treatment with immune checkpoint blockade has only been reported in retrospective studies[28]. As the therapeutic landscape of metastatic RCC rapidly evolves, the role of cytoreductive nephrectomy in the immunotherapy era is currently evaluated in randomized clinical trials (PROBE, NORDIC-SUN trials)[29]. Here, we show that tumor cell killing rate is significantly higher with CD8 T cells isolated post-nephrectomy compared to baseline. In line with this, we observed that radical nephrectomy is associated with elevation of peripheral tumor-reactive CD8 T cells expressing the effector markers CX3CR1 and GZMB in a majority of patients and regardless the metastatic status. Our data corroborates with a prior study showing a significant reduction in circulating exhausted CD8 T cells (BTLA^+^) between baseline and 30 days post nephrectomy[30]. Altogether, our data suggest that some patients may benefit from radical nephrectomy to eliminate immunosuppressive signals (e.g soluble PD-L1) generated by the primary tumor and restore a pool of circulating tumor-reactive CD8 T cells with high cytotoxic activity.

Predicting response to immune checkpoint blockade prior to initiation of treatment or early in the course of treatment remains a critical unmet need as it will help reduce overtreatment, treatment-related toxicities and healthcare expenditures while improving patient outcome. In addition, selecting the right treatment among the available immunotherapies is a challenging scenario faced by patients and clinicians. To overcome this challenge, we established an ex-vivo individualized functional assay that quantitatively measures the cytotoxic activity of CD8 T cells isolated from patient peripheral blood. We tested the efficacy of FDA-approved immune checkpoint inhibitors (pembrolizumab, nivolumab and atezolizumab) on the killing activity of CD8 T cells isolated from RCC patients post-nephrectomy. Most patients demonstrated improved T-cell mediated killing activity upon PBMC treatment with immune checkpoint inhibitors including those with no detectable killing activity at baseline. In all cases, atezolizumab treatment led to inferior induction of T-cell killing activity compared to other immune checkpoint inhibitors which corroborates with the limited activity of atezolizumab as adjuvant and front-line therapy in RCC[31]. T-cell killing rates were not statistically different between pembrolizumab and nivolumab but divergence in response to treatment was observed at the individual patient level. Out of 13 patients, 5 patients showed an increase of more than 20% in killing rates with nivolumab compared to pembrolizumab. Inversely, only 1 patient showed similar increase with pembrolizumab compared to nivolumab. At this time, it is unknown whether nivolumab can lead to superior efficacy compared to pembrolizumab as no head-to-head comparison studies have been conducted yet. Pembrolizumab has been recently approved for adjuvant treatment in RCC[32] but primary results for nivolumab are not expected before 2024. Both antibodies bind PD-1 with very high affinity and specificity and their pharmacological characteristics are similar. Despite they bind the same target, differential response to pembrolizumab and nivolumab for the same patient suggest distinct but uncharacterized yet mechanism(s) of action.

Immune checkpoint blockade has significantly improved overall survival in patients with advanced cancers but only a minority of patients experience complete and durable response. Novel immunotherapies are still needed for patients with resistant and refractory advanced cancers. In line with this, our group has recently developed a novel monoclonal antibody, called H1A, that binds PD-L1 and downregulates its expression at the plasma membrane[9]. Cell-surface expression of PD-L1 is regulated by CMTM6 (CKLF Like MARVEL Transmembrane Domain Containing 6) which protects PD-L1 from lysosomal and proteasomal degradation[33, 34]. H1A binding to PD-L1 reduces PD-L1/CMTM6 interaction leading to PD-L1 destabilization and lysosomal degradation[9]. The unique mechanism of action of H1A can kill two birds with one stone as PD-L1 degradation can not only prevent the extrinsic function of PD-L1 (i.e PD-L1/PD-1 binding) but also inhibit pro-tumorigenic PD-L1 intracellular signaling[35]. In this study, we showed that treatment of PBMC with H1A boosts T-cell mediated killing activity. Noteworthy, we found a significant increase in T-cell killing rates with H1A compared to pembrolizumab and nivolumab supporting the therapeutic potential of H1A antibody as next-generation immunotherapy.

Atezolizumab and H1A bind the same target (i.e PD-L1) but have distinct mechanisms (PD-L1/PD-1 blockade versus PD-L1 degradation). To investigate the underpinnings of the superior antitumor activity of H1A, we compared the phenotype of PBMC treated with atezolizumab and H1A by mass cytometry. Surprisingly, in atezolizumab-treated group, we observed an enrichment in regulatory T cells expressing immunosuppressive molecules PD-1 and LAG-3 which can explain the inefficient tumor cell killing ability of CD8 T cells observed *in vitro*. It has been previously reported that PD-1/PD-L1 blockade induces proliferation and tumor infiltration of PD-1^+^ regulatory T cells leading to poor patient response to immunotherapy and in some cases hyperprogressive disease[36, 37]. Targeting regulatory T cells in combination with PD-1/PD-L1 therapy has shown promise in preclinical studies and reports from clinical studies are still awaited [38-40]. LAG-3 is an immune checkpoint expressed by regulatory T cells which stimulates the release of immunosuppressive cytokines IL-10 and TGF-β dampening the antitumor immune response[41]. Combination of LAG-3 inhibitors with PD-1/PD-L1 blockade is evaluated in clinical trials and it has already showed positive results in melanoma[42, 43]. As opposed to atezolizumab, RCC patient-derived PBMC treated with H1A displayed accumulation of proliferative CD8 T and NK cells with an effector phenotype (Granzyme B^+^T-bet^+^) and reduced levels of regulatory T cells. Our immunophenotyping data corroborate with the robust cytotoxic killing of tumor cells observed *in vitro*.

We recognize few limitations in our study. All patients enrolled in the study did not receive immune checkpoint inhibitors following nephrectomy, therefore we were not able to assess the predictive value of our ex-vivo functional assay with patient clinical response. With the recent approval of immune checkpoint inhibitors as front-line treatment for metastatic RCC and adjuvant treatment for localized high risk RCC, such work will be feasible in a near future. The size of the study cohort is small and further validation is warranted using a larger number of patients. Cytotoxic activity of CD8 T cells were measured with an immortalized RCC tumor cell line (786-O) as reference target cell line to assess the impact of PD-L1/PD-1 blockade in the immune cell compartment. Future work using patient-derived tumor cells is ongoing to determine the translational potential of our ex-vivo functional assay in evaluating response to immune checkpoint inhibitors in an individualized manner.

## Conclusions

The low rates of durable responses to immune checkpoint inhibitors in RCC stress the need to develop tools to predict patient response to immunotherapy and design novel and effective immunotherapies. In this proof-of-concept, we showed differential T-cell cytotoxic activity in response to various FDA-approved immune checkpoint inhibitors. Finally, we demonstrated the therapeutic value of a newly developed PD-L1 monoclonal antibody that induces proliferation and cytotoxic activity of RCC patient-derived CD8 T and NK cells.

## Acknowledgments

This work was supported by NIH R01CA256927 (HD) and R01AI095239 (HD). FL was recipient of a postdoctoral fellowship from the Fonds de Recherche du Quebec-Sante (FRQS). We thank generous patients who have participated to the study and provided blood samples.

## Author Contributions

HD and FL conceptualized the study. ZA, MAH, JKG, SMH, HZ, TX, JBH performed and analyzed *in vitro* experiments. KDP performed mass cytometry experiments and Rphenograph analysis. CL, RN, RRP, VS, HT and BCL collected demographics, clinical characteristics and supervised the clinical study protocol. FL wrote the first draft of the manuscript. All the authors reviewed and approved the manuscript.

## Disclosures

A patent application has been filed with Mayo Clinic regarding the development and application of the H1A antibody. No conflict of interest is reported by all authors.

---

**Supplementary Figure 1:**
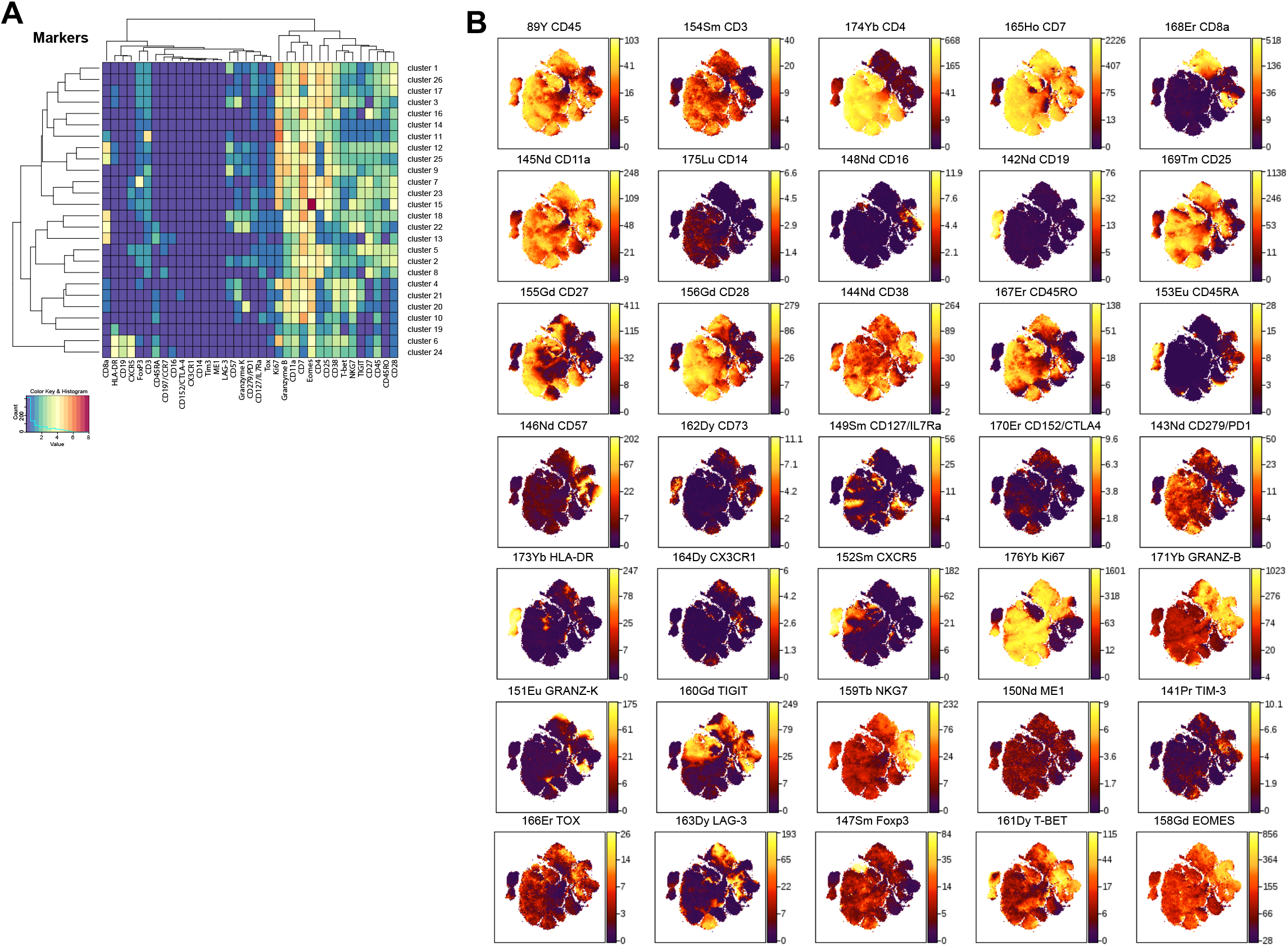
Clustering of RCC patient-derived PBMC analyzed by mass cytometry. A) Heatmap demonstrating the distribution and relative intensity of 36 immune cell markers used in the clustering analysis. B) Density plots of 26 immune cell markers on patient-derived PBMC analyzed by mass cytometry

## Notes

### Competing Interest Statement

The authors have declared no competing interest.

### Summary of Updates

New data have been provided in regards to the immunophenotyping of PBMC treated with H1A and atezolizumab

